# Dynamic mode structure of active turbulence

**DOI:** 10.1101/2022.04.15.488501

**Authors:** Richard J. Henshaw, Olivia G. Martin, Jeffrey S. Guasto

## Abstract

Dense suspensions of swimming bacteria exhibit chaotic flow patterns that promote the mixing and transport of resources and signalling chemicals within cell colonies. While the importance of active turbulence is widely recognized, the structure and dynamics of the resulting collective flows are the subject of intense investigation. Here, we combine microfluidic experiments with proper orthogonal decomposition (POD) analysis to quantify the dynamical flow structure of this model active matter system under a variety of conditions. In isotropic bulk turbulence, the modal representation shows that the most energetic flow structures dictate the spatio-temporal dynamics across a range of suspension activity levels. In confined geometries, POD analysis illustrates the role of boundary interactions for the transition to bacterial turbulence, and it quantifies the evolution of coherent active structures in externally applied flows. Beyond establishing the physical flow structures underpinning the complex dynamics of bacterial turbulence, the low-dimensional representation afforded by this modal analysis offers a potential path toward data-driven modelling of active turbulence.

## INTRODUCTION

The collective motion of concentrated suspensions of swimming bacteria and self-propelled actin filaments are typical of a broad class of active systems, ranging from flocking birds and schooling fish down to micron-sized synthetic active colloids [1–5]. Such active suspensions regulate fundamental ecological and biological processes for example in the gut microbiome of the human gastrointestinal tract [6], as well as the mitotic spindle structure during cell division [7]. At high concentrations, swimming bacteria form emergent jet-like flow structures and eddies [8–12], which are correlated over length scales that are one order of magnitude larger than the individual cells and enhance the colony mobility [13]. These chaotic flow patterns – termed as active turbulence [14] – are reminiscent of classical, inertial turbulence [15] and exhibit a range of novel physical properties, such as: enhanced material transport [16], boundary-driven pattern formation [17, 18], and reduced viscosity and super-fluidlike behavior [19, 20]. Apart from traditional, bulk statistical measures and velocity spectra [3, 4, 11, 12], relatively little progress has been made in understanding the underlying flow structure of active turbulence, especially in the presence of boundaries and ambient flows. A host of theoretical and computational models have been proposed to explain these phenomena [4, 21–23], as well as the onset of active turbulence under confinement [24], and recent work using machine learning approaches to uncover the equations of motion for active turbulence are promising [25]. However, the vast experimental parameter space and necessity for big data sets to characterize these chaotic systems complicate data-driven discovery through simulations. Thus, establishing new approaches to characterize the dynamic structure of active turbulence and provide compact reduced-order representations are essential, for example to understand the movement ecology of collective microbial motion [26] and to design and engineer new classes of active materials [27].

Inspired by approaches from inertial turbulence, we apply proper orthogonal decomposition (POD) to analyse the spatio-temporal dynamics of self-generated bacterial turbulence flows. POD – also known as PCA or the Karhunen-Loève procedure [28–30] – computes an orthogonal basis set that optimally represents the fluid velocity field, an approach that has been instrumental for example in the development of reduced-order flow models [29, 31, 32]. In POD, a velocity field **u**(**x**, *t*) at *N* discrete times is represented by a linear combination of *N* orthonormal spatial modes, *ϕ*_*n*_(**x**), as

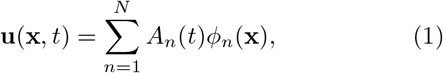

where 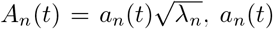 is a normalized temporal coefficient, and *λ*_*n*_ is the kinetic energy for a given mode [32]. This process is achieved through singular value decomposition (SVD; *Materials and Methods*) [33], which ranks the spatial modes, *ϕ*_*n*_(**x**), by decreasing energy, *λ*_*n*_. Despite the striking similarities between inertial and active turbulence, POD techniques have seen limited applications soft matter, almost exclusively for granular flows [34–36], with low Reynolds number flows typically overlooked due to their generally linear nature.

In this work, we use microfluidic experiments to measure the chaotic self-generated flow fields of bacterial turbulence under a variety of conditions, which we then decompose using modal analysis to quantify their flow structure and dynamics. Bulk bacterial turbulence is firstly investigated far from lateral boundaries, where we determine the effects of varying cell activity [4, 12] on the spatio-temporal mode structure. Subsequently, we use POD to elucidate the role of boundary interactions in the transition to bacterial turbulence in confined geometries [18, 24], and to quantify the evolution of coherent active structures in externally applied flows [37]. Not only does POD analysis unveil the constituent flow structures underpinning the chaotic bacterial motion in each of these physical systems, but this framework also provides an accurate and efficient representation of active turbulence with remarkably few spatial modes.

## RESULTS AND DISCUSSION

### Modal analysis of bulk bacterial turbulence

In unbounded domains, concentrated suspensions of motile bacteria exhibit complex spatio-temporal flow patterns. Dense suspensions of motile *B. subtilis* bacteria were loaded into microfluidic chambers (26 *μ*m deep) with widely separated side walls (2 mm). The active suspensions were imaged far from the boundaries using video microscopy for ≈1 min (*N* = 6300 frames), and time-dependent velocity fields, **u**(**x**, *t*), characterizing the bacterial motion (Fig 1*A*) were measured using particle image velocimetry (PIV; *Materials and Methods*) [18]. The resulting chaotic flows exhibit vortex structures (≈40 *μ*m) and mean flow speeds (⟨|**u**|⟩ ≈50 *μ*m.s^−1^) that are significantly larger than the respective length and swimming speed of individual cells, consistent with previous observations [11, 12].

**FIG. 1.**
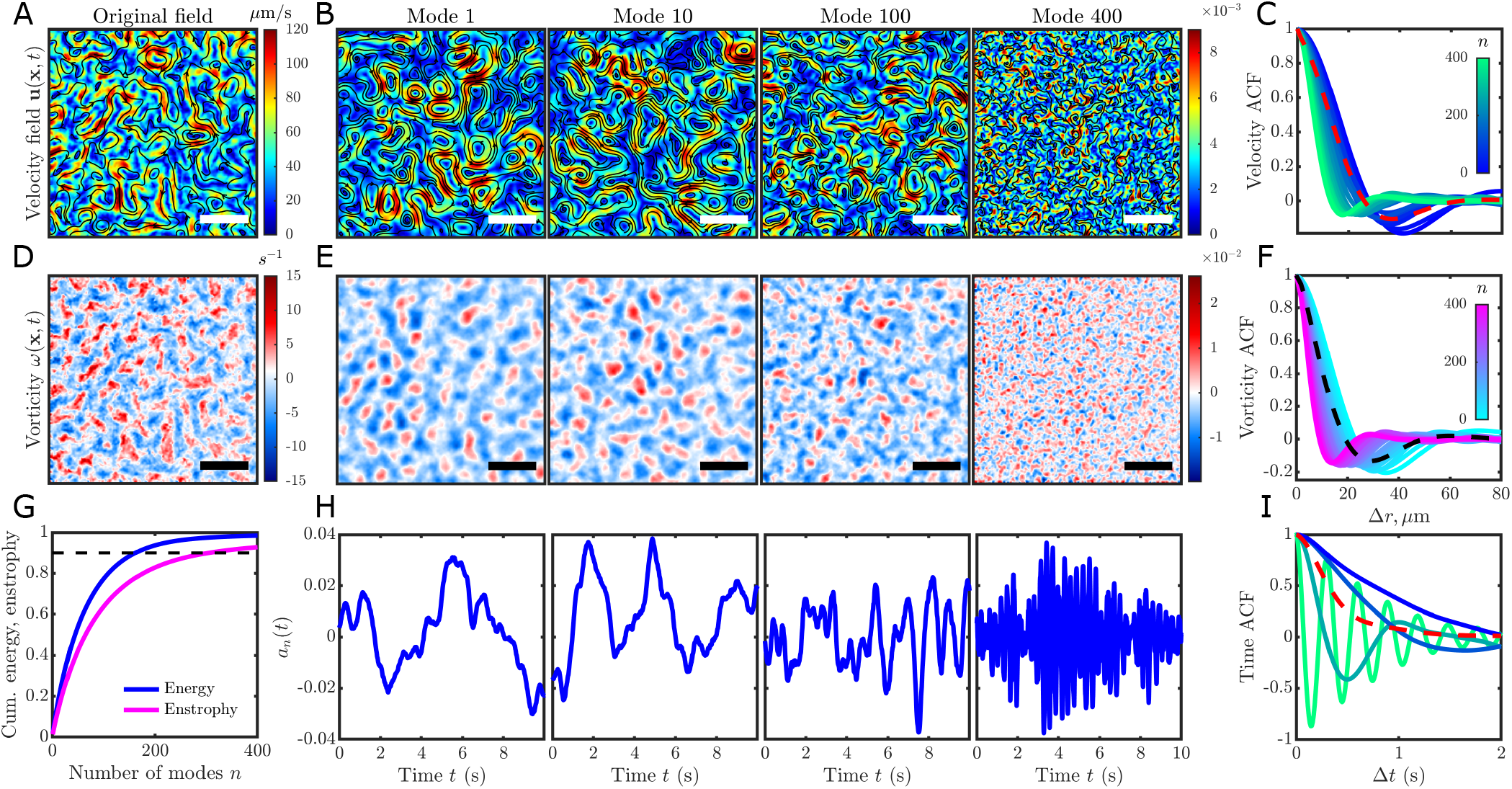
POD reveals dominant spatio-temporal structures of bacterial turbulence. (*A*) Instantaneous measured velocity field, **u**(**x**, *t*), obtained from the collective motion of a dense suspension of swimming bacteria. (*B*) Sample velocity field POD modes, *ϕ*_*n*_(**x**), organized by decreasing kinetic energy. (*C*) Spatial autocorrelation function (ACF) of the first 400 POD velocity field modes compared to the original flow field (dashed red), showing a decrease in characteristic length scale as the mode number increases. (*D*) Instantaneous vorticity field, *ω* = **∇** × **u**, corresponding to (*A*). (*E*) Sample vorticity field POD modes organized by decreasing enstrophy. (*F*) ACF of POD vorticity field modes compared to the original vorticity field (dashed black). (*G*) Normalized cumulative mode energy (blue) and enstrophy (magenta) 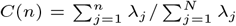 for the first *n* modes. 162 and 305 modes are required to capture 90% (black dashed) of the cumulative energy and enstrophy, respectively. (*H*) Temporal amplitudes, *a*_*n*_(*t*), of the velocity field modes in (*B*). (*I*) Time ACF for the mode amplitudes shown in (*H*) compared to the velocity field time ACF (dashed red). Colors correspond to (*C*). Scale bars are 100 *μ*m.

Energy-ranked velocity field modes from POD analysis reveal the hierarchy of flow structures underlying bacterial turbulence, whereby the highest energy modes are typically associated with the largest structures. The dominant flow structures are contained within the first several modes (Fig. 1*B, n* = 1, 10, 100), whilst the higherorder modes capture minor perturbations to the flow field (Fig. 1*B, n* = 400) as well as measurement noise. The autocorrelation function (ACF) of the spatial modes (Fig. 1*C*) illustrates a cascade from large to small flow structures with decreasing mode energy (increasing *n*). Furthermore, the highest energy modes (*n* ≲ 100) encompass the dominant spatial scales of the original velocity field (Fig. 1*C*, dashed red).

Analogous to the velocity field, a POD analysis of the vorticity field (Fig 1D), *ω* = ∇ × **u**, provides an enstrophy-ranked set of vorticity modes (Fig. 1*E*). The vorticity mode ACFs clearly illustrate a similar refinement of the modal vortex size (Fig. 1*F*) with decreasing enstrophy. Considering the cumulative information contents (Fig. 1*G*), only *N*_*KL*_ = 162 modes (2.5%) and 305 modes (4.8%) are required to capture 90% of the cumulative energy and enstrophy, respectively, based on the Karhunen-Loève (KL) dimension [38]. The larger number of modes for the latter suggests that enstrophy is more equally distributed across flow structures compared to the kinetic energy, although this effect may be exacerbated by numerical gradients used to calculate the enstrophy.

Beyond the spatial structure, modal analysis provides insight into the temporal dynamics of the chaotic flow fields. The temporal coefficients, *a*_*n*_(*t*), describe the instantaneous amplitude (Fig. 1*H*) for each spatial mode (Fig. 1*B*) and fluctuate strongly in time. The ACFs of the temporal coefficients (Fig. 1*I*) show that the large, high energy flow structures associated with low mode numbers are strongly correlated. While lower energy modes exhibit faster initial decay rates, the envelope of their rapidly fluctuating ACF is comparable to correlation strengths of the high energy modes and the original velocity field (Fig. 1*I*, red dashed). Thus, not only does POD provide a rigorous framework for concise low-order representations of this model active matter system, but it also establishes new tools to gain deeper insight into their complex structure and dynamics under a variety of physical conditions.

### Effect of bacterial activity on mode structure and energy distribution

Suspension activity is a critical factor in determining the dynamics of active turbulence [39]. Bacterial activity is quantified by the mean flow speed of the suspension, ⟨|**u**|⟩, and varied here in a controlled manner by exploiting the aerobic properties of the bacteria [11]. Microfluidic chambers were loaded with cells as described previously, and the bacterial suspensions were imaged periodically over ≈30 min. As oxygen was slowly depleted, the cell activity decreased from > 50 *μ*m.s^−1^ to approximately the single-cell swimming speed (≈15 *μ*m.s^−1^). For all activities, the kinetic energy spectra, *E*(*k*), as a function of the wave number, *k*, collapse (Fig. 2*A*) to previously reported power-law behaviors [4], illustrating the robustness of these results.

**FIG. 2.**
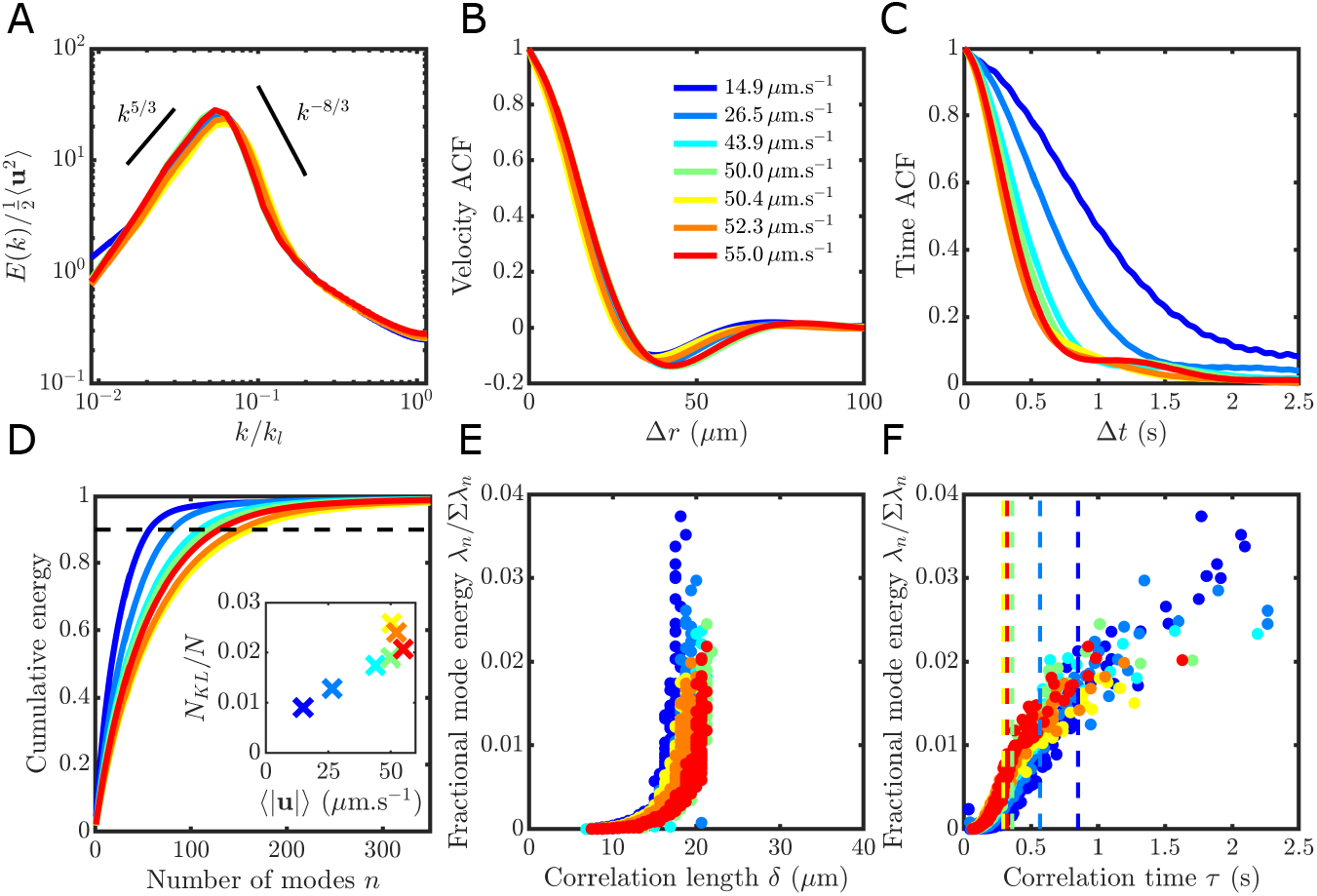
Variation in bacterial turbulence structure with swimming activity. (*A*) Normalized kinetic energy spectra as a function of normalized wave number exhibit self-similarity and consistent power-law behavior [4] across a wide range of bacterial activity levels, ⟨|**u**|⟩. The characteristic wave number is *k*_*l*_ = 2*π/l*, and *l* = 4 *μ*m is the typical cell length. (*B*) Spatial velocity ACF remains constant across various cell activities. (*C*) Temporal velocity ACF broadens significantly as bacterial swimming activity decreases. (*D*) Cumulative kinetic energy contained within POD modes illustrates the relative increase in energy for low order modes with decreasing cell activity. Black dashed line corresponds to the 90% energy KL dimension. (*E*) Fractional mode energy as a function of the correlation length, *δ*, of individual POD modes (*n* ≤ 400; 1*/e* decay of velocity ACF; Fig. 1*C*) show that the most energetic modes typically correspond to the largest spatial scales, which remain constant across bacterial activities. (*F*) Fractional mode energy as a function of the correlation time, *τ*, of individual POD mode amplitudes (*n* ≤ 400; 1*/e* decay of time ACF; Fig. 1*I*). The most energetic modes have the largest *τ*, which increase with decreasing activity. Vertical dashed lines indicate *τ* for velocity field from (*C*).

Beyond classical measures of collective motion, modal analysis reveals rich changes in the structure and complexity of bacterial turbulence with cell activity, including a diminishing number of dominant modes as the cell swimming speed decreases. Across all activities, the characteristic length scale of the turbulent velocity fluctuations remains approximately constant [11, 12], as shown by the spatial velocity ACF (Fig. 2*B*). Consequently, the slowdown results in a broadening of the temporal ACF (Fig. 2*C*). However, the underlying changes in the spatial and temporal dynamics are more comprehensively captured by POD analysis. As the cell activity decreases, the energy distribution shifts to lower order modes (Fig. 2*D*). For example, the lowest activity examined (⟨|**u**|⟩ = 14.9 *μ*m.s^−1^) required 65% fewer modes to capture 90% of the kinetic energy relative to the most active suspensions (Fig. 2*D*, inset). Moreover, analysis of the spectra of mode correlation lengths, *δ*, reveals that dominant spatial modes emerge (Fig. 2*E*), which are characterized by both high energy and long correlation lengths and reflect *δ* for the original flow field (Fig. 2*B*). While the spectra of *δ* exhibit only a minor shift toward lower correlation lengths, the correlation times of the mode amplitudes, *τ*, increase strongly with decreasing cell activity (Fig. 2*F*). This analysis illustrates that the most energetic velocity field modes are also responsible for the increase in *τ*. Taken together, these analyses show how the spatio-temporal complexity of bacterial turbulence markedly decreases with decreasing cell activity.

### Edge modes illustrate the role of confinement in the transition to active turbulence

Boundaries play a fundamental role in regulating active turbulence [39], whereby confinement has been shown to select for vortical flow structures [17] and lead to the formation of self-organized flow patterns [40]. Similar to inertial turbulence, bacterial suspensions [18, 41, 42] and active nematics [24, 41, 43, 44] confined in a channel exhibit a striking transition from stable flow to chaotic dynamics as the separation between lateral boundaries increases. By replicating recent experimental results (Fig. 3*A-H*) [18], we extend our POD analysis to quantify the effects of confinement on bacterial turbulence. Briefly, a dense suspension of *B. subtilis* was confined laterally in a series of microfluidic “racetrack” channels (see *Materials and Methods*) with varying width, *W* (Fig. 3*A*). Strong confinements – where *W* is approximately less than the bulk vortex size (Fig. 1) – rectify cell activity into a unidirectional flow (Fig. 3*C*), as shown by the stream-wise averaged flow profile (*F* (*y*); Fig. 3*F*) and normalized net flow rate (*ψ*; Fig. 3*B*; see *Materials and Methods*). As *W* increases, vortex structures emerge (Fig. 3*D*) before giving way to chaotic dynamics (Fig. 3*E*), when *W* is significantly larger than the bulk vortex size. The formation of vortexes and the subsequent chaotic flow are typified by sinusoidal velocity profiles (Fig. 3*G-H*) with strong wall velocities and a rapid decrease in *ψ* (Fig. 3*B*), both of which are in excellent agreement with previous work [18]. Reduced order POD representations accurately and compactly capture the emergent active velocity field behaviors across varying channel widths (Fig. 3*F-H*) throughout this flow transition.

**FIG. 3.**
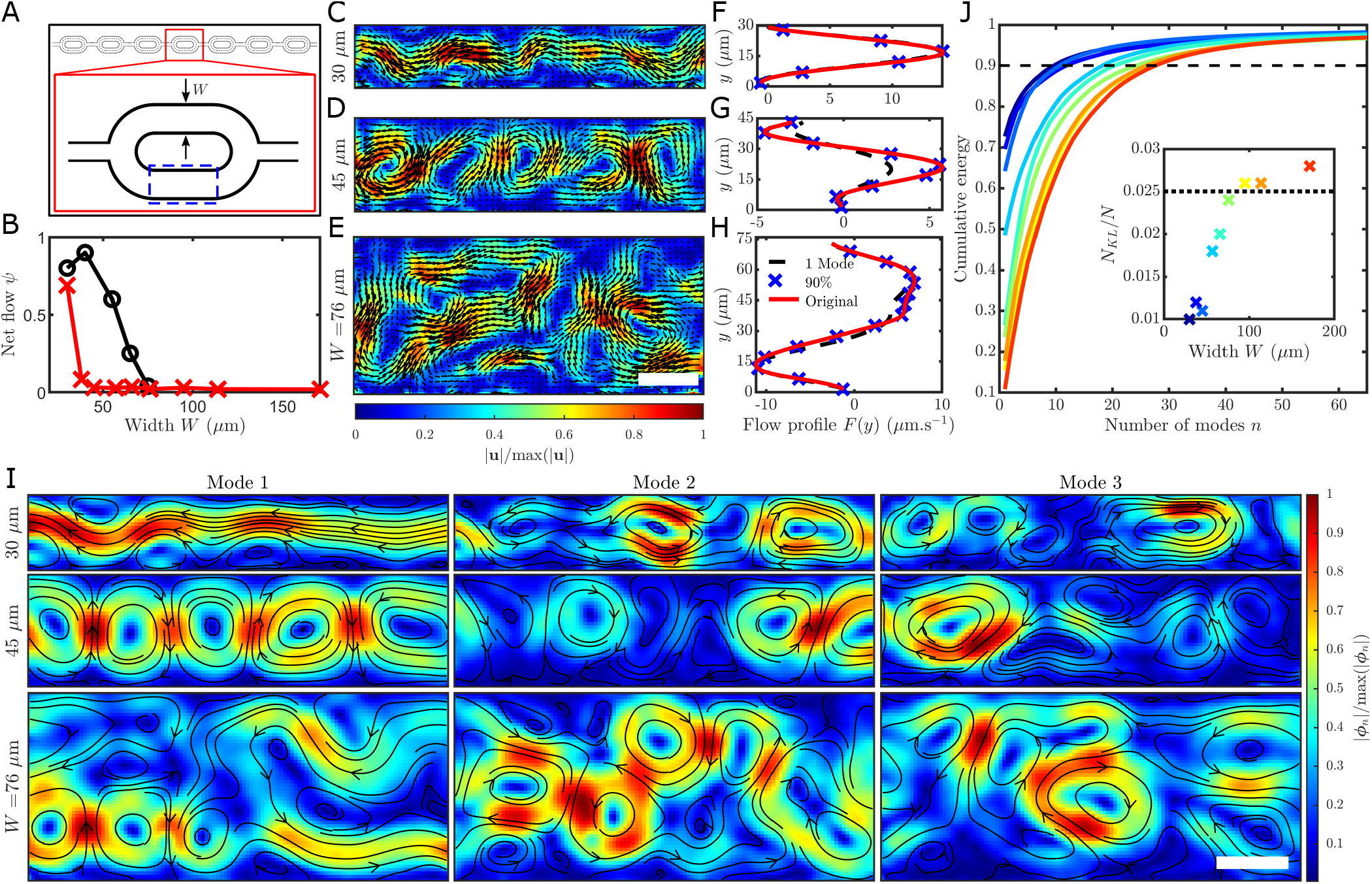
POD unveils the role of edge modes in the transition to active turbulence under confinement. (*A*) Schematic of microfluidic racetrack channels of varying widths (30 *μ*m ≤ *W* ≤ 171 *μ*m) to study effects of confinement on bacterial turbulence [18]. Straight interrogation regions (dashed blue) were fixed at ≈ 150 *μ*m long. (*B*) Normalized net flow rate, *ψ*, illustrates the transition from unidirectional flow for small *W* to turbulent flow at large *W* in the current study (red crosses), which compares well to previous work (black circles) [18]. (*C-E*) Normalized instantaneous velocity fields for various channel widths. Scale bar, 25 *μ*m. (*F-H*) Stream-wise averaged flow profiles, *F* (*y*), for original flow fields (red; corresponding to (*C-E*)) compared to a 90% POD reconstruction (blue crosses) and single mode reconstruction (black dashed) using modes from (*I*). (*I*) The three highest energy (normalized) POD modes for various *W*. Scale bar, 25 *μ*m. (*J*) Cumulative POD mode energy increases with decreasing channel width. Black dashed line indicates the 90% energy KL dimension. (Inset) *N*_*KL*_ as a fraction of the total number of modes, *N*, plateaus as the channel approaches bulk conditions (black dotted).

The topology of the spatial modes, *ϕ*_*n*_(**x**), reveal the fundamental flow structures underpinning the effects of confinement on suspension dynamics (Fig. 3*I*). For the strongest confinement examined, the highest energy mode encapsulates the nearly-steady, stable flow (Fig. 3*I, W* = 30 *μ*m), representing > 70% of the total kinetic energy (Fig. 3*J*). At intermediate channel widths, mode 1 highlights the emergent vortex chain, where higher-order modes have similar topologies that describe stream-wise fluctuations of the vortexes (Fig. 3*I, W* = 45 *μ*m). Importantly, in weak confinement prior to transitioning to isotropic, bulk turbulence, the most energetic mode reveals strong, wall-driven activity (mode 1; Fig. 3*I, W* = 76 *μ*m). Such edge modes are consistent with both previous experimental observations of confined bacterial suspensions [18] and the pinning and stabilization of defects to walls in active nematics simulations [39], which are thought to regulate the transition to active turbulence. In contrast, lower energy modes (modes 2-3) describe chaotic fluctuations in the center of the channel away from boundaries, akin to bulk turbulence modes (Fig. 1*B*). The kinetic energy distribution across modes is a strong indicator of the onset of bacterial turbulence under confinement (Fig. 3*J*), which is exemplified by the rapid drop of the kinetic energy in mode 1 from 70% to 10% as *W* increases from 30 *μ*m to 171 *μ*m. Moreover, the number of modes, *N*_*KL*_, to capture 90% of the system energy exhibits a striking increase with *W* (Fig. 3*J*, inset) that plateaus at *W* = 95 *μ*m (or 2.5 vortex lengths in the bulk). The fractional *N*_*KL*_ required at large *W* is consistent with bulk turbulence results (Fig. 3*J*, inset, dashed; see also Fig. 1) and indicates that the system has reached the fully chaotic regime.

### Evolution of active turbulence structures in externally driven flow

Driven ambient flows are known to strongly modify the dynamics of active suspensions [37, 45, 46], leading to novel rheological [19, 20] and transport [37] properties. Pressure-driven flows of dense, swimming bacterial suspensions through microfluidic channels were recently observed to exhibit significant flow intermittency [37]. Active vortex structures interact with the imposed external flow to produce sporadic periods of near-parabolic and plug-like flow in motile suspensions of *B. subtilis* (Fig. 4*A*), compared to non-motile cells with steady flow (Fig. 4*E*). Here, we investigate this phenomenon through a POD analysis of previously published data [37]. The most energetic velocity field mode for the motile (Fig. 4*B*) and non-motile (Fig. 4*F*) suspensions are both representative of the parabolic background flow. For the active suspension, vortex chain structures (Fig. 4*C-D*) characterize higher-order modes (*n* ≥ 2), responsible for the observed flow intermittency. In contrast, the featureless higher-order modes for the non-motile cells (Fig. 4*G-H*) comprise a comparatively minute fraction of the total kinetic energy (Fig. 4*I*), commensurate with the nature of their steady, homogeneous flow. The non-trivial higher-order modes of the swimming bacterial suspension also give direct insights into the dynamics of the active flow structures. The vortex chain modes, *ϕ*_2_(**x**) and *ϕ*_3_(**x**), both appear periodic in space with a wavelength, ℓ = 320 *μ*m, and a cross-correlation analysis reveals a phase difference of Δ*x* = 84.5 *μ*m (Fig. 4*K*), which is consistent with the orthogonal nature of POD modes. Similarly, the temporal coefficients (Fig. 4*J-K*), *a*_2_(*t*) and *a*_3_(*t*), exhibit a period and phase difference *T* = 1.27 s and Δ*t* = 0.333 s, respectively. Because propagating turbulent flow structures can be characterized by a linear combination of (minimum) two modes, we can estimate their speed via a phase velocity as *v*_*P*_ = ℓ */T* ≈ 252 *μ*m.s^−1^. This result is comparable to the flow structure velocity (≈235 *μ*m.s^−1^) measured by manual tracking (*Materials and Methods*) [37]. A principle application of POD is the identification of coherent structures in flow [47–50], which is effectively demonstrated here in the case of active turbulence.

**FIG. 4.**
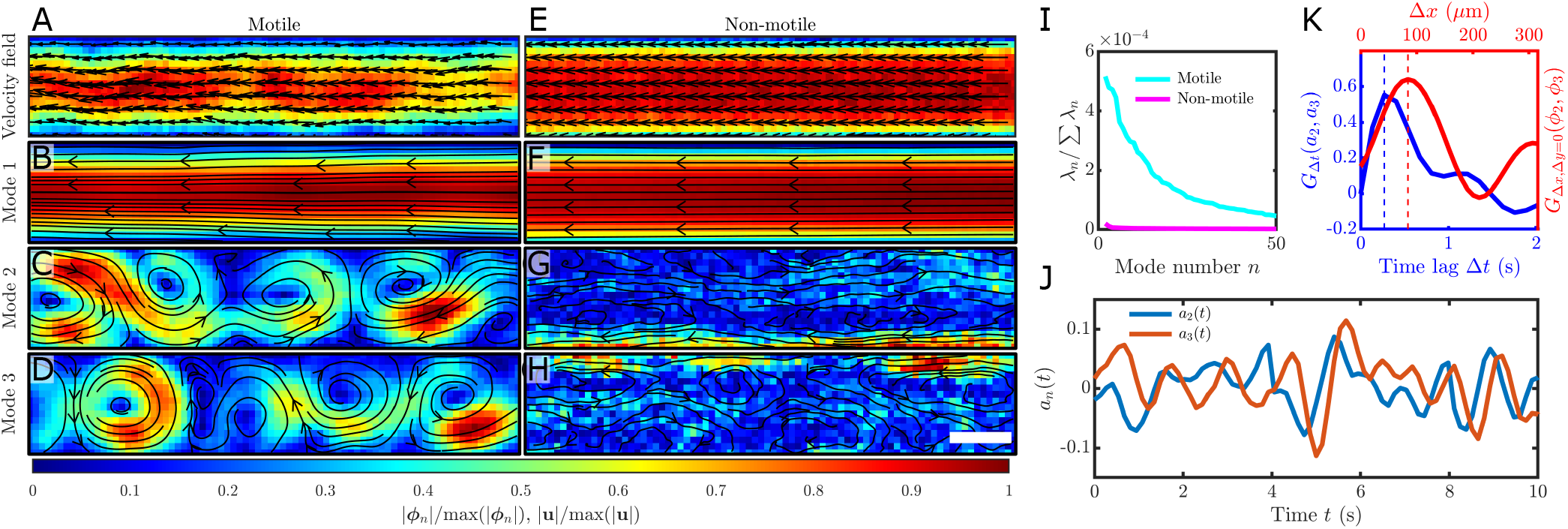
POD modes capture intermittent bacterial turbulence in externally applied flows. (*A*) Instantaneous normalized velocity field of a dense, active bacterial suspension flowed through a rectangular microchannel exhibits strong flow intermittency. Velocity field data obtained from previously published work [37]. (*B-D*) Three most energetic (normalized) POD modes for the motile suspension in (*A*). Mode 1 resembles a Poiseuille-like background flow, whereas modes 2 and 3 are anti-phase vortexes that capture travelling active flow structures. (*E*) Instantaneous normalized velocity field of a non-motile cell suspension under the same conditions as (*A*), which shows a near-parabolic flow profile. (*F-H*) Three most energetic (normalized) POD modes for the non-motile suspension in (*E*). Mode 1 resembles a Poiseuille-like background flow, while modes 2 and 3 capture only minor noise fluctuations. Scale bar, 100 *μ*m. (*I*) Fractional POD mode energy for the motile (cyan) and non-motile (magenta) suspensions corresponding to (*A-D*) and (*E-H*), respectively. (*J*) Temporal amplitudes *a*_2_ (blue) and *a*_3_ (orange) of the motile suspension, corresponding to spatial modes 2 and 3 in (*C*) and (*D*), respectively. (*K*) Phase shifts of the temporal mode amplitudes (blue) and spatial modes (red) for modes 2 and 3 are indicative of the traveling turbulent structures shown in (*A*). *G*_Δ*t*_ is the temporal cross-correlation of mode amplitudes shown in (*J*), and *G*_Δ*x*,Δ*y*=0_ is the spatial cross-correlation in the stream-wise direction of POD modes shown in (*C-D*).

## CONCLUSION

In this work, we established the application of proper orthogonal decomposition to gain new physical insights into the collective motion of dense suspensions of swimming bacteria. Far from boundaries, POD analysis of bulk bacterial turbulence revealed that the spatial and temporal structure of the underlying velocity field modes strongly correlated with kinetic energy, which naturally ranks the mode structures in decreasing size. Decreasing cell activity shifts energy to lower-order modes that are responsible for increasing the correlation time [4, 12], effects that are masked when considering only the bulk field. For confined cell suspensions, our results accurately capture the transition to chaotic flow [18], and the resulting velocity field modes illustrate the importance of boundary interactions in regulating the dynamics, as predicted by active nematics simulations [24]. In external flows, higher-order velocity field modes describe the propagation of coherent active flow structures that are responsible for the observed flow intermittency [37]. Despite their prevalence in inertial turbulence and macroscale flow applications [32], the potential insights gained from POD and related techniques in microscopic flows have largely been overlooked, due to the relatively predictable behavior of low Reynolds number flows. However, as illustrated here, the chaotic nature of active matter flows presents copious opportunities for mode analysis techniques to yield new insights into the dynamics and potentially the stability [51] of these complex systems. As demonstrated by our present work, POD provided an efficient, compact description of experimental data for collective bacterial motility. Such low dimensional representations of active turbulence will facilitate new data driven models of these phenomena [52, 53], for example through machine learning techniques [25].

## MATERIALS AND METHODS

### Cell culturing

Wild-type *Bacillus subtilis* bacteria (strain OI1084) were taken from −80°C frozen stock and streaked onto 1.5% agar plates prepared with Terrific Broth (TB, Sigma). Plates were incubated at 25°C for 24 hours, after which time a single colony from the plate was used to inoculate an overnight liquid TB culture at 30°C with shaking (200 rpm). The bacterial suspension was then subcultured (1.5 ml of cell culture into 60 ml of pre-warmed TB) and grown at 35°C and 200 rpm for 6 hours to mid-log phase (OD_600_ ≈ 0.2). Immediately prior to experiments, dense cell suspensions (∼ 10^10^ cells.ml^−1^) were prepared by centrifugation at 5000 g for 5 minutes, and the pellet was resuspended with 2 *μ*l of fresh TB media.

### Microfluidics and image analysis

Polydimethlysiloxane (PDMS) microfluidic channels were fabricated through soft lithography [54] and plasma bonded to No. 1 thickness glass coverslips. The PDMS channels were thinly cast to ensure ample diffusion of oxygen to the bacterial suspension and prolonged cell activity. Dense cell suspensions were gently loaded into the microfluidic devices via pipette, and the channel inlet and outlet were sealed with wax to prevent residual flows. For all experiments, bacterial suspensions were imaged with brightfield illumination on an inverted microscope (Nikon Ti-E) using a sCMOS camera (Zyla 5.5, Andor Technology). Time-resolved velocity fields, **u**(**x**, *t*), of the bacterial suspensions were measured by performing Particle Image Velocimetry (PIV) using PIVLab [55] implemented in MATLAB. The subsequent velocity fields were then lightly smoothed using a Guassian kernel with a standard deviation of one PIV pixel (1.73 *μ*m) in space and one frame (9.5 ms) in time. Vorticity fields, *ω*(**x**, *t*), were computed from measured velocity fields using central differencing (MATLAB). The POD was performed on the resulting velocity and vorticity fields using built-in singular value decomposition functions in MATLAB.

### Bulk cell suspensions with varying activity

To capture the bulk dynamics of dense bacterial suspensions in the absence of lateral wall effects (Fig 1-2), large microfluidic chambers (2 mm ×2 mm ×26 *μ*m) were prepared, where the side length corresponds to ≈80 correlation lengths of the turbulent bacterial suspension. Imaging was performed with a 20 × objective (0.45 NA) at 105.5 fps. Bacterial activity (Fig. 2) was varied using a previously established approach [11, 12]. Briefly, the cell suspensions were imaged periodically over the course of 30 minutes, during which time the cell swimming speed naturally decayed due to oxygen depletion, where the decay rate of cell activity was controlled via the thickness of the PDMS device. Without the need to manipulate the cell suspension, this approach ensured that the bacterial concentration was constant across varying activity levels. Seven data sets were captured in total, which consisted of 6, 300 frames each (≈ 1 min per video) and a 2 − 3 min delay between acquisitions. Data analysis was restricted to the central portion of the chamber, ≈ 30 correlation lengths from the lateral boundaries.

### Cell suspensions under varying confinement

To quantify POD modes for bacterial suspensions under varying degrees of confinement (Fig. 3), experiments from [18] were replicated with minor modifications. Microfluidic devices (19 *μ*m deep) were fabricated, which comprised a series of racetrack geometries connected by narrow inlets (30*μ*m wide). Nine racetrack widths were chosen between 30 *μ*m ≤ *W* ≤171 *μ*m in fractional increments of our measured characteristic vortex size (38 *μ*m) taken as the minimum point in the spatial ACF in bulk conditions (Fig. 1C, dotted red line). Cell suspensions were imaged at 40× (0.6 NA) magnification and 100 fps for 10 s per channel width. All confined microfluidic geometries were simultaneously loaded with the same cell suspension and imaged within the first 10 min of loading to ensure consistent bacterial activity. Experiments were repeated across multiple days with freshly cultured bacteria to verify repeatability. The net flow *ψ* was calculated following [18] as *ψ* = | (Σ**u ê**_*x*_) */* (Σ ||**u**||)|, where **ê**_*x*_ is the unit vector along the principal channel direction and the sum is over all PIV sub-windows over 5 s (500 frames) of video. *ψ* = 1 indicates unidirection circulation flow around the racetrack, and *ψ* = 0 corresponds to a globally stationary suspension. The stream-wise flow profiles *F* (*y*) were generated from the velocity field by averaging over time and space as *F* (*y*) = ⟨**u**(*x, y, t*)· **ê**_*x*_⟩ _(*x,t*)_, where **ê**_*x*_ was chosen such that *F* (*y*) is on average positive.

### Cell suspensions under externally imposed flow

The data corresponding to Fig. 4 was obtained from a previously published work [37], where the relevant experimental details are repeated here for convenience. Dense suspensions (≈1.3–2 × 10^10^ cells.ml^−1^) of wild-type *B. subtilis* (strain OI1085) were flowed through a PDMS microfluidic chamber (200 *μ*m wide × 320 *μ*m deep) via a syringe pump at a flow rate of *Q* = 25 *μ*l.h^−1^. Imaging was performed under brightfield conditions at 30× magnification. For motile cell suspensions, 19 individual videos (4 s duration) were acquired at 370 fps. Velocity fields were quantified through ghost particle velocimetry [56] and averaged such that each pixel is the average of 32 × 32 pixel square over 50 frames. Image data and velocity fields for non-motile cell suspensions were acquired under the same conditions, but with a dead suspension of *B. subtilis*. Vortical regions in the flow were identified through use of the Okubo-Weiss parameter [57, 58] for manual tracking of travelling flow structures.

## ACKNOWLEDGMENTS

We thank Eleonora Secchi for generously providing the data used in Fig. 4, and Michael Stehnach for assistance with microfluidic device fabrication for Fig. 3. We also thank Hugo Wioland, Jörn Dunkel, Sami Yamanidouzisorkhabi, and Irmgard Bischofberger for valuable discussions, and Louison Thorens for comments on the manuscript. This work was supported by NSF awards CAREER-1554905 and OCE-1829827.

